# Olfactory Response as a Marker for Alzheimer’s Disease: Evidence from Perception and Frontal Oscillation Coherence Deficit

**DOI:** 10.1101/840934

**Authors:** Mohammad Javad Sedghizadeh, Hadi Hojjati, Kiana Ezzatdoost, Hamid Aghajan, Zahra Vahabi, Heliya Tarighatnia

## Abstract

High-frequency oscillations of the frontal cortex are involved in functions of the brain that fuse processed data from different sensory modules or bind them with elements stored in the memory. These oscillations also provide inhibitory connections to neural circuits that perform lower-level processes. Deficit in the performance of these oscillations has been examined as a marker for Alzheimer’s disease (AD). Additionally, the neurodegenerative processes associated with AD, such as the deposition of amyloid-beta plaques, do not occur in a spatially homogeneous fashion and progress more prominently in the medial temporal lobe in the early stages of the disease. This region of the brain contains neural circuitry involved in olfactory perception. Several studies have suggested that olfactory deficit can be used as a marker for early diagnosis of AD. A quantitative assessment of the performance of the olfactory system can hence serve as a potential biomarker for Alzheimer’s disease, offering a relatively convenient and inexpensive diagnosis method. This study examines the decline in the perception of olfactory stimuli and the deficit in the performance of high-frequency frontal oscillations in response to olfactory stimulation as markers for AD. Two measurement modalities are employed for assessing the olfactory performance: 1) An interactive smell identification test is used to sample the response to a sizable variety of odorants, and 2) Electrophysiological data are collected in an olfactory perception task with a pair of selected odorants in order to assess the connectivity of frontal cortex regions. Statistical analysis methods are used to assess the significance of selected features extracted from the recorded modalities as Alzheimer’s biomarkers. Olfactory decline regressed to age in both healthy and AD groups are evaluated, and single- and multi-modal classifiers are also developed.

The novel aspects of this study include: 1) Combining EEG response to olfactory stimulation with behavioral assessment of olfactory perception as a marker of AD, 2) Identification of odorants most significantly affected in AD patients, 3) Identification of odorants which are still adequately perceived by AD patients, 4) Analysis of the decline in the spatial coherence of different oscillatory bands in response to olfactory stimulation, 5) Being the first study to quantitatively assess the performance of olfactory decline due to aging and AD in the Iranian population.

## Introduction

Alzheimer’s disease (AD) is the most prevalent type of dementia affecting approximately one individual in 10 in the population older than 65 (1). Early diagnosis of AD is necessary to ensure that the required clinical and social care are provided for affected individuals (2). AD is known to be associated with the aggregated deposition of surplus amyloid-beta (Aβ) protein, a product of synaptic activity (3), as plaques causing neurotoxic events such as inflammation and synaptic loss, and other neural degenerations linked to another protein, phosphorylated tau (4).

Accumulated levels of these proteins in the brain have been measured as biomarkers for AD through PET imaging of the brain or sampling the cerebrospinal fluid (CSF) (5) (6). Several neuropsychological tests have also been introduced to evaluate the mental state of subjects in a clinical exam. Mini-Mental State Exam (MMSE) and Mini-Cog are two examples of such tests, which are used by clinicians to evaluate the cognitive skills of the patients and decide upon further evaluation tests (7) (8) (9). Although no single protocol has been established for large-scale screening of AD, there are proposed frameworks for diagnosis based on a set of biomarkers such as those just mentioned (10). An update to the National Institute on Aging - Alzheimer’s Association (NIA-AA) Research Framework provides additional flexibility for introducing new biomarkers to allow the results of new measurement modality evaluations in observational studies to establish their value in the clinical assessment of AD (11).

In addition to the mentioned assessment methods, previous studies have shown that olfactory deficit is an early symptom of Alzheimer’s disease (12) (13). Further studies have demonstrated that standard methods of assessing olfactory system such as sniffing kits can be helpful in distinguishing AD patients from healthy individuals (14) (15) (16).

The neurodegenerative processes associated with the deposition of neurofibrillary tangles and amyloid plaques in the brain do not progress in a homogeneous fashion (17), and are more prominent in the medial temporal lobe in the early stage of the disease (18). Interestingly, the medial temporal lobe is the region where olfactory perception also occurs. Therefore, perception of smells is affected more severely in AD patients compared to its decline caused by normal aging, and several studies have suggested that olfactory deficit can be used as a biomarker for early diagnosis of AD (19) (20).

Unlike PET imaging or CSF sampling, measuring the odor perception abilities of patients is an inexpensive and non-invasive procedure. However, the perception of odors highly correlates with the culture of the individuals, and hence, the familiarity of subjects with the employed odorants needs to be considered in defining a smell scoring procedure. The University of Pennsylvania smell identification test (UPSIT) (21) proposes a standard sniffing kit for assessing the olfactory function and has been used for studying the olfactory deficit in AD patients (22). However, some of the odors used in this test may be unfamiliar for non-American societies. To address this issue, researchers have proposed alternative scents in modified sniffing kits to create tests that are suitable for their populations of interest (23) (24) (25).

A practical issue in administering sniffing kit tests is that they require the participant’s cooperation. Patients with dementia-like symptoms may have difficulty following the written test questions or the clinician’s instructions or may respond erroneously due to not recalling the name of an odorant they indeed perceived. Also, the way an examiner interacts with the participant may introduce bias towards specific options in the response sheet [34]. To circumvent the interference of non-olfactory related issues in the performance of the tests, methods relying on EEG recording during the presence of odorants to participants have been proposed. One such technique is the olfactory event-related potential (OERP), in which a sequence of odorants is presented to the participant at regular time instances, allowing the EEG response data to be averaged over several trial intervals for reducing noise and enhancing the fidelity of the recorded data. Several studies have focused on the role of OERP test results as an early biomarker for AD. In (26) OERP waveforms analyzed and spotted features for differentiating AD patients and an age-matched control group. In a more recent study (27), OERP was employed to distinguish between AD and mild cognitive impairment (MCI) patients.

Another method for the differential analysis of EEG data of AD patients and healthy participants is coherence analysis. Coherence refers to the functional connectivity of different brain regions and is measured by the synchrony of oscillations recorded at different EEG electrodes. Earlier studies have indicated that the coherence of EEG channels can help in the diagnosis of AD (28) (29) (30). These studies showed that the reduction in the functional connectivity of the brain regions is captured as a decrease in the coherence between EEG channels (31). Some reports have also assessed the relative value of the coherence of EEG channels for different frequency bands in the classification of AD patients and healthy participants (32) (33).

In this paper, we examine the characteristics of olfactory response as markers for the diagnosis of AD through the use of both EEG and behavioral olfactory response data. Our EEG analysis comprises an assessment of the statistical significance of the coherence in the EEG data across the spatial domain for different frequency bands. In the behavioral olfactory response data, our approach identifies the best subset of odorants among those in a localized version of the UPSIT kit (Iran-SIT (16)), which significantly contributes to classifying AD patients from healthy participants. MMSE scores are also used as reference for evaluating our olfactory-based results. Single-modality regressors are developed employing the significant components identified in each set of olfactory response data (EEG coherence and behavioral UPSIT) separately. The regressors are age-adjusted to account for the decline in the performance caused by normal aging. Furthermore, by employing the statistically significant components from both modalities, we propose a multi-modal classifier of AD patients versus healthy participants, which also regresses the olfactory decline due to aging.

## Materials and Methods

Figure 1 illustrates an overview of the multi-modal data analysis methodology used in our study.

**Figure 1.**
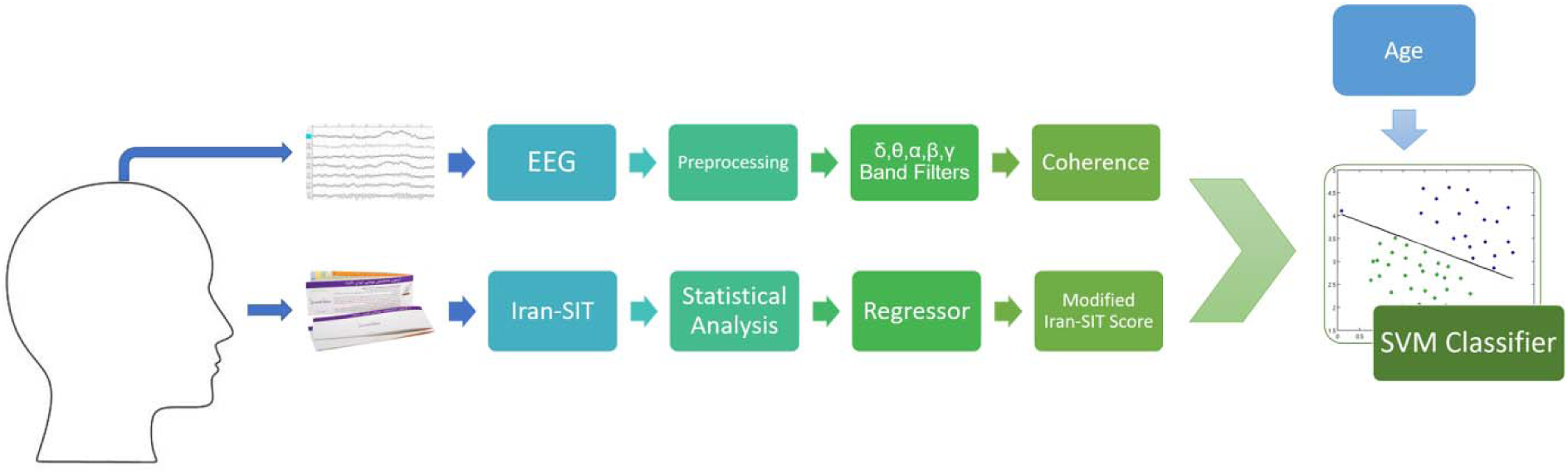
An overview of our methodology. The coherence between EEG electrode pairs in different frequency bands, the modified Iran-SIT score, and the age of participants are used as features for training an SVM classifier. The modification of the Iran-SIT score as well as the selection of significant frequency bands and connections in the EEG records are carried out by statistical analysis.

### Participants

This study was approved by the Review Board of Tehran University of Medical Sciences and all participants gave their written consent to participate in the experiment. Participants were selected among the individuals referring to the memory clinics of two hospitals in Tehran (Ziaeian Hospital and Roozbeh Hospital, both affiliated with Tehran University of Medical Sciences) with memory performance complaints. All the tests were carried out in the Department of Geriatric Medicine of Ziaeian Hospital in South of Tehran. Two expert neuropsychologists assessed all the participants and recorded their smoking history, preferred hand, age, level of education, as well as any past olfactory problem. Demographic and medical history data were also collected for each participant. A total of 52 participants were recruited for the study, and after applying exclusion criteria (as described in the next subsection), twenty-four individuals (age = 72.1 ± 9.0, female = 54.25%), including 11 participants with AD (age = 76.6 ± 9.2, female = 64%) and 13 healthy participants (age = 68.2 ± 6.2, female = 46%) were selected for data analysis. The Mini-Mental State Examination (MMSE), the Clock Drawing Test (CDT), and a verbal fluency test were performed. After the neuropsychological assessment, a neurologist examined the participants and conducted the Functional Assessment Scales Test (FAST) (34).

Then, the participants performed the UPSIT examination and after a few minutes of rest, performed the EEG-based olfactory measurement test. Table 1 shows the overall statistics of the participants. Details of the clinical assessment procedure and the MMSE, UPSIT, and EEG-based experiments are described in the following subsections.

**Table 1.**
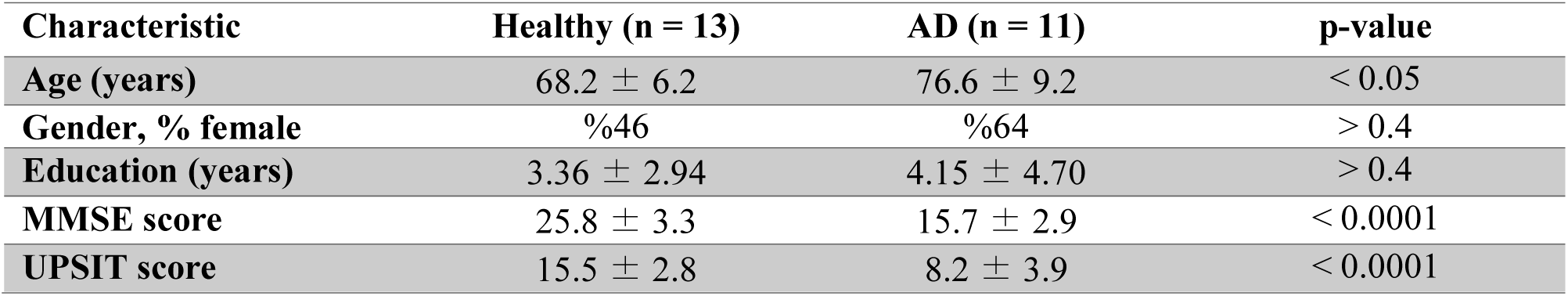
Participant characteristics. p-values denote the separation between healthy participants and AD patients in each characteristic.

| Characteristic | Healthy (n = 13) | AD (n = 11) | p-value |
| --- | --- | --- | --- |
| Age (years) | $68.2 \pm 6.2$ | $76.6 \pm 9.2$ | $< 0.05$ |
| Gender, % female | %46 | %64 | $> 0.4$ |
| Education (years) | $3.36 \pm 2.94$ | $4.15 \pm 4.70$ | $> 0.4$ |
| MMSE score | $25.8 \pm 3.3$ | $15.7 \pm 2.9$ | $< 0.0001$ |
| UPSIT score | $15.5 \pm 2.8$ | $8.2 \pm 3.9$ | $< 0.0001$ |

### Clinical Diagnostic Assessment

An expert neurologist diagnosed probable Alzheimer’s disease according to the latest guideline of the NIA-AA (35). AD participants must meet the criteria prescribed for diagnosing dementia as described in (35). Results of the Mini-Mental State Examination (MMSE) and inquiry about the onset and progressions of the symptoms from the patients and their companions were used by the neurologist to diagnose cognitive impairment. In addition to criteria for dementia, AD participants must also meet the criteria for probable AD dementia. Structural MRI images (1.5 Tesla MR Scanner and a 16-channel HR head coil) were analyzed, and the Medial Temporal Atrophy Scale, White Matter Lesions, and Global Atrophy Scale were used for describing the image. Exclusion criteria were a history of stroke, schizophrenia, major depressive disorders and electroconvulsive therapy (ECT) over the past six months, traumatic brain injury, non-AD neurodegenerative diseases (Parkinson’s disease, Progressive Supranuclear Palsy, Multi-System Atrophy, Cortico-Basal Degeneration), and any history of olfactory pathway disorders. MCI patients were also excluded from this study.

### Mini-Mental State Examination (MMSE)

MMSE (36) is a clinical test commonly used for measuring cognitive impairment. During the MMSE test, different memory skills are evaluated, and a score out of 30 is produced. Based on this score and the education level of the patients, clinicians assess the participant’s cognitive skills.

MMSE consists of the following cognitive test categories: Orientation to time, Orientation to place, Registration, Attention and calculation, Delayed recall, Naming, Repetition, Reading, Writing, Visio-spatial, and Commands. A score is given in each category, and the sum of a participant’s scores in all categories is used as the MMSE score. It should be mentioned that due to the low literacy levels of many of the participants in this study, lower MMSE scores were registered in both AD and healthy control groups.

### University of Pennsylvania Smell Identification Test (UPSIT)

UPSIT and its modified versions have been employed in earlier studies for early diagnosis of AD. Due to the culture-specific nature of smell perception, the reliability of these tests has to be evaluated in different populations (2) (37). Localized versions of the UPSIT test kit have been introduced in countries such as Brazil (23), Turkey (24), Lithuania (25), and Iran (16) (for an exhaustive review refer to (14)).

To this date, no research results based on the UPSIT kit or other olfactory-based tests have been reported for the detection of AD in the Iranian population. The current study is the first to utilize a localized version of the UPSIT test to diagnose AD in its early stages in Iran.

The test kit consists of 24 odors which are each exposed by scratching its corresponding strip. The list of the odors is included in supplementary materials. After presenting each scent to the participant, four options to select from are provided, and the participant is asked to identify the closest match among these options to the odor that they perceived. As some participants in the study were not able to read the list of options printed in the kit, either because of vision problems or due to illiteracy, the list of options for each odor presentation was read loudly and clearly to the participants, once before and once after the presentation of each odor.

#### Classification

To assess the results of the UPSIT test, we employed a support vector machine (SVM) classifier with a linear kernel to separate AD patients from healthy participants based on the UPSIT score and the age of the participants. Normal aging is known to be a major cause for olfactory deficit and hence, when dealing with the UPSIT or other olfactory test results, it is essential to take into account the effect of aging.

Due to the small size of our dataset, we used 5-fold cross-validation to evaluate the accuracy of the classifier. In this evaluation scheme, the dataset is divided into five equal folds, and each time, the label of one fold is predicted using the model trained on data of the other folds.

#### Statistical Analysis of UPSIT

The UPSIT score denotes the number of correct answers for each participant. However, among the twenty-four odors of the test, some are probably more effective at separating Alzheimer’s patients from healthy participants. To determine these significant odors, each participant’s answers were converted to a vector of binary elements in which zeros represent wrong answers and ones indicate correct answers. These vectors are divided into two groups of AD patients and healthy participants and then a welch’s t-test (as well as one-way ANOVA test) is applied to samples from these groups. By doing this, twenty-four p-values corresponding to the presented odors were obtained. The Benjamini and Hochberg method was applied for p-value correction (38). The remaining small p-values (p-value < 0.05) indicate odors that are significant in separating the two participant groups. The Scipy and Statsmodels packages were used for statistical analysis and p-values < 0.05 were considered as significant.

After identifying the significant odors, we performed a linear regression analysis to obtain a modified UPSIT score as a weighted sum of the significant odor scores. The coefficient of each score in the linear regressor is set to maximize the separability of the AD patients and healthy participants.

We fitted a linear regression model to each of the AD patients and healthy participant groups using the modified UPSIT score and age. These lines demonstrate how the olfactory perception weakens with age in the AD and healthy groups. Having a participant’s modified UPSIT score and considering their age, the regressor model classifies the participant into one of the two groups. For comparison, we also derived similar age-regressed lines using the raw UPSIT score.

#### EEG-based Olfactory Measurement

EEG signals were recorded using a 32-channel Mitsar amplifier. Data were recorded from the Fz, Cz, Pz, and Fp1 electrodes. The Fz, Cz, and Pz channels were chosen based on the results of a similar study (27). The Fp1 electrode data was used to identify eye movements and eye blinks. The choice of using a limited number of channels in this study was to accommodate for the age and mental condition of many of the participants, so the time to install and confirm the functionality of the electrodes would be kept to a minimum. The channel impedance was maintained under 15 kΩ for each electrode. The EEG sampling rate was 2000 Hz, and electrodes were referenced to the A1 earlobe (39). EEGLAB was used for data preprocessing (40).

Participants performed an olfactory perception task (41). During this task, the participant is presented with a sequence of stimuli composed of two different odors, one of which occurring more frequently (standard) and the other being presented rarely (deviant) (42). Lemon was chosen as the frequent stimulus and Rose as the rare one (27). Our experimental protocol consisted of a two-second stimulus presentation followed by 8 seconds of rest (pure water) interval. The odors were delivered to the participants using a laboratory olfactometer (43). The probability of rare stimuli was 0.25 (27). Each trial (epoch) took 10 seconds, and the whole experiment for each participant consisted of 120 trials and took about 20 minutes. The 90 frequent and 30 rare odors were presented in random but preset order.

The choice of odors in olfactory experiments is an important issue. When the objective is to test the performance of the olfactory system, odors must be selected so as not to arouse the trigeminal system. This is because the olfactory and the trigeminal systems are interconnected and may interact by intensifying and suppressing each other during exposure to certain stimuli (44). Therefore, in our experiments, we replaced the eucalyptus odor which was used in other studies (27) with lemon, since eucalyptus excites both the olfactory and trigeminal systems. We also increased the duration of odor presentation to two seconds to allow for regular breathing cycles by the participants (to accommodate for their age and mental condition). The extracted event epochs included one second of pre-stimulus and two seconds of post-stimulus data, following the empirical estimate of the olfactory response latency of about 600-700 milliseconds (45) (46).

#### EEG Preprocessing

Signals were filtered to 0.5 to 40.5 Hz and downsampled to 200 Hz. Independent Component Analysis (FastICA (47)) was used for eye blink removal. Then, EEG signals were divided into epochs based on the stimulus onset, and heavily artifact-contaminated epochs were manually excluded.

#### Coherence Analysis

EEG data were filtered using the delta (0.5-3.99 Hz), theta (4.0-7.99 Hz), alpha (8.0-12.99 Hz), beta (13.0-29.99 Hz), and gamma (30.0-40.5 Hz) filters to separate the different oscillatory bands. The gamma filter had an upper limit of 40.5 Hz since the EEG recording system was programmed to remove the higher frequency components, including the 50 Hz powerline interference. Coherence analysis was employed at frequent (lemon) epochs for each frequency band. For each pair of connections between the four channels, the imaginary part of coherence was measured for each frequency band using Eq. (1):

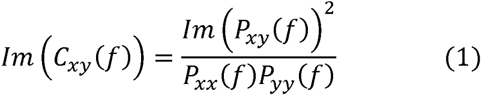

in which *P_xy_* (*f*) indicates the cross power spectral density between channels *x* and *y* for frequency band *f*, and *P_xx_* (*f*) indicates the power spectral density of channel x for frequency band *f*.

A total of 30 coherence values (5 bands times six connections) were calculated. The Welch t-test (38) was applied, and p-values for all the coherence values were calculated. Due to the redundancy in the coherence values (data from the six connections have some degree of correlation), we used the Benjamini and Hochberg correction method using an effective sample size calculation method described in Chapter 8 of (38). Coherence values with p-value <0.05 were considered significant.

To assess the performance of the resulting significant coherences in classifying AD patients and healthy participants, an SVM classifier was trained using the significant coherences and age as its features.

#### Multimodal Analysis

An objective of this study is to propose a multi-modal method for distinguishing AD patients from healthy participants based on the two olfactory-based data types. This is to suggest the most efficient combination of olfactory-based tests for diagnosing AD. To achieve this goal, an SVM classifier with a linear kernel was used to separate AD patients from healthy participants based on the significant components of the modified UPSIT test and the significant coherence values between the EEG electrodes calculated in different frequency bands. The age of the participants was also considered as a feature in training the classifier. A 5-fold cross-validation was used to evaluate the accuracy of the classifier.

## Results

### MMSE

Details of the MMSE results are shown in Table 2. This table includes the participant’s age, diagnosis state, total MMSE score as well as the MMSE score in each category. The p-value for each test category is also shown.

**Table 2.** MMSE results. Abbreviations used: Edu = Education (years), O Time=Orientation to Time, O Place=Orientation to Place, Reg=Registration, AttCalc=Attention and Calculation, DelRec=Delayed Recall, Name=Naming, Rep=Repetition, Read=Reading, Write=Writing, VisSpat=Visuo-Spatial, Comm=Commands. Data of exluded participants are not shown in the table.

| Subject | Diagnosis | Age | Edu | MMS | O Time | O Place | Reg | AttCal | DelRe | Name | Rep | Read | Writ | VisSpa | Comm |
| --- | --- | --- | --- | --- | --- | --- | --- | --- | --- | --- | --- | --- | --- | --- | --- |
| 1 | AD | 75-80 | 6 | 19 | 4 | 4 | 3 | 0 | 0 | 2 | 1 | 1 | 1 | 1 | 2 |
| 2 | Normal | 75-80 | 6 | 27 | 5 | 5 | 3 | 4 | 2 | 2 | 1 | 1 | 0 | 1 | 3 |
| 3 | AD | 75-80 | 0 | 16 | 2 | 5 | 3 | 0 | 2 | 2 | 1 | 0 | 0 | 0 | 1 |
| 4 | AD | 80-85 | 3 | 18 | 4 | 5 | 3 | 1 | 0 | 2 | 0 | 0 | 0 | 0 | 3 |
| 5 | Normal | 70-75 | 5 | 30 | 5 | 5 | 3 | 5 | 3 | 2 | 1 | 1 | 1 | 1 | 3 |
| 6 | AD | 85-90 | 0 | 12 | 2 | 3 | 3 | 0 | 0 | 2 | 1 | 0 | 0 | 0 | 2 |
| 7 | AD | 70-75 | 4 | 15 | 2 | 5 | 3 | 0 | 0 | 2 | 0 | 0 | 0 | 0 | 3 |
| 8 | AD | 85-90 | 0 | 16 | 3 | 4 | 3 | 0 | 0 | 2 | 1 | 0 | 0 | 0 | 3 |
| 9 | Normal | 65-70 | 6 | 26 | 5 | 5 | 3 | 4 | 2 | 2 | 0 | 1 | 0 | 1 | 3 |
| 10 | Normal | 65-70 | 12 | 30 | 5 | 5 | 3 | 5 | 3 | 2 | 1 | 1 | 1 | 1 | 3 |
| 11 | Normal | 70-75 | 0 | 21 | 5 | 5 | 3 | 1 | 2 | 2 | 1 | 0 | 0 | 0 | 3 |
| 12 | Normal | 60-65 | 14 | 29 | 5 | 5 | 3 | 5 | 2 | 2 | 1 | 1 | 1 | 1 | 3 |
| 14 | Normal | 65-70 | 0 | 21 | 5 | 5 | 3 | 1 | 1 | 2 | 1 | 0 | 0 | 0 | 3 |
| 15 | AD | 65-70 | 5 | 12 | 3 | 2 | 3 | 0 | 0 | 2 | 0 | 0 | 0 | 0 | 2 |
| 16 | Normal | 55-60 | 4 | 27 | 5 | 5 | 3 | 2 | 3 | 2 | 1 | 1 | 1 | 1 | 3 |
| 17 | AD | 65-70 | 0 | 18 | 2 | 3 | 3 | 0 | 3 | 2 | 1 | 0 | 0 | 0 | 3 |
| 19 | AD | 70-75 | 6 | 19 | 5 | 3 | 3 | 3 | 0 | 2 | 0 | 1 | 1 | 0 | 1 |
| 20 | AD | 80-85 | 8 | 17 | 2 | 5 | 3 | 0 | 0 | 2 | 0 | 1 | 1 | 0 | 3 |
| 21 | AD | 60-65 | 5 | 11 | 3 | 2 | 3 | 0 | 0 | 2 | 0 | 0 | 0 | 0 | 3 |
| 22 | Normal | 75-80 | 0 | 23 | 4 | 5 | 3 | 4 | 2 | 2 | 1 | 0 | 0 | 0 | 3 |
| 23 | Normal | 70-75 | 0 | 26 | 5 | 5 | 3 | 5 | 1 | 2 | 1 | 1 | 0 | 0 | 3 |
| 24 | Normal | 70-75 | 6 | 26 | 5 | 4 | 3 | 2 | 3 | 2 | 1 | 1 | 1 | 1 | 3 |
| 25 | Normal | 65-70 | 0 | 21 | 5 | 4 | 3 | 1 | 3 | 2 | 0 | 0 | 0 | 0 | 3 |
| 26 | Normal | 60-65 | 12 | 28 | 5 | 5 | 3 | 3 | 3 | 2 | 1 | 1 | 1 | 1 | 3 |
|  | p-value |  |  |  | <0.05 | <0.05 | >0.05 | <0.05 | <0.05 | >0.05 | <0.05 | <0.05 | >0.05 | <0.05 | <0.05 |

### UPSIT

An SVM classifier was trained to classify the AD and healthy participants using the total UPSIT score and age as its features. The accuracy of the SVM classifier using 5-fold cross-validation was 83.3% with an area under the curve (AUC) of 0.87. These results suggest that there is a relationship between age, olfactory functionality (represented by the UPSIT score) and the diagnosis of AD.

As shown in Figure 2a, the olfactory functionality decreases with age in both the AD and healthy groups. However, the decline is more significant in the AD group and can be distinguished from the deficit caused by normal aging.

**Figure 2.**
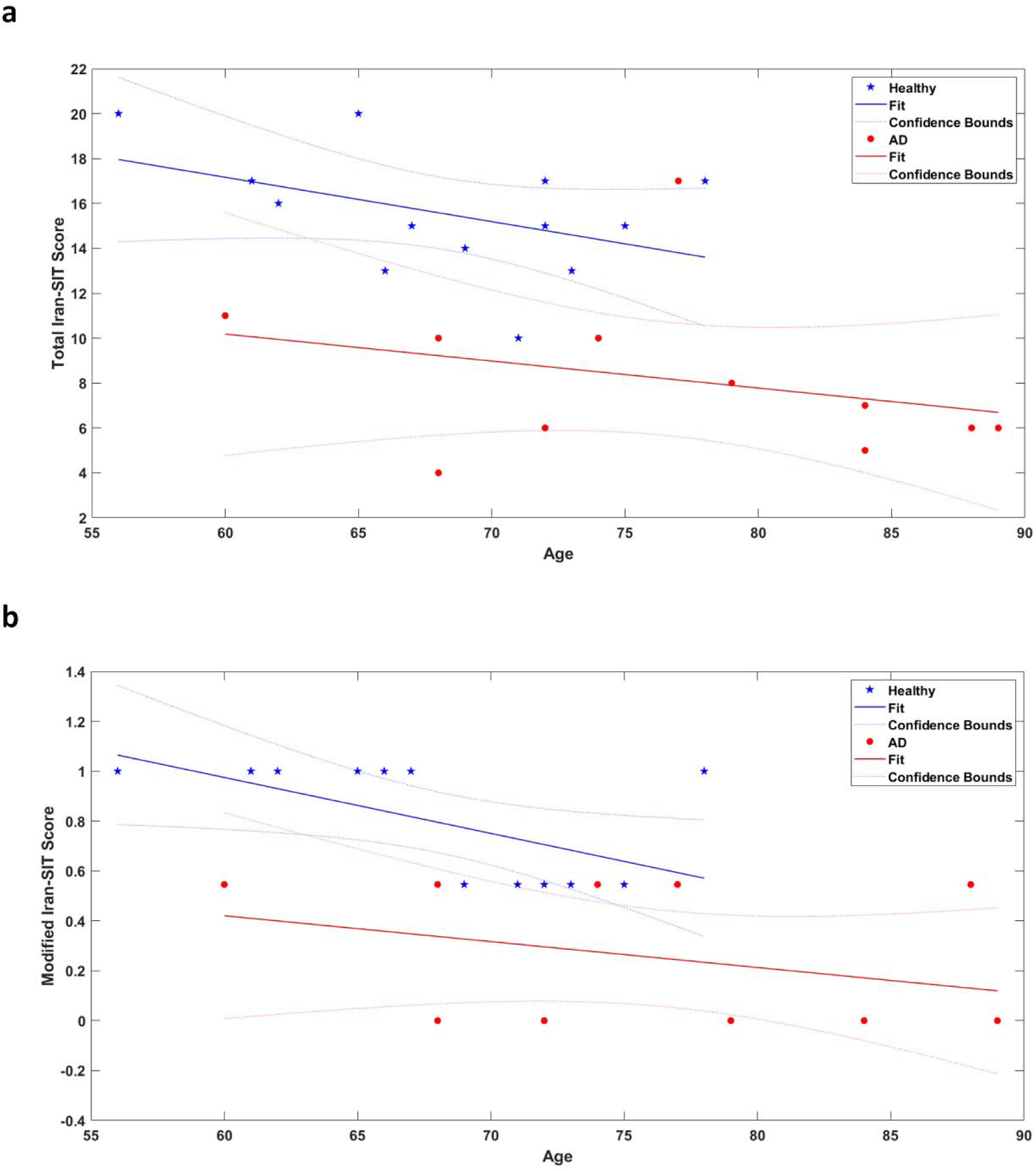
The total and modified UPSIT (Iran-SIT) scores plotted versus age: a) Total UPSIT score, b) Modified UPSIT score. A regression line is fitted to the data of each group in both figures, and the confidence bounds are also plotted.

As mentioned in the previous section, p-value was calculated for each odor to identify the most sensitive odorants. Two significant odors (p-value < 0.05) were identified (Grape and Chocolate). These odors, as well as their p-values, are shown in Table 3.

**Table 3.** p-values derived from the ANOVA one-way test for significant odors separating the two groups of participants.

| Odorant | p-value |
| --- | --- |
| Q6 (Grape) | <0 .05 |
| Q21 (Chocolate) | <0 .05 |

We then calculated the modified UPSIT score for each participant by summing the responses to the two sensitive odors. The resulting modified score versus age is plotted in Figure 2b. It can be seen that by only using the two significant odors, the test can separate AD patients from healthy participants.

The MMSE and total UPSIT scores of AD patients and healthy participants are plotted versus age in a three-dimensional plot in Figure 3a. It can be seen that the two groups are relatively separable. However, there are still several data points that cannot be correctly classified.

In order to analyze whether our modifications can cause complete separability of the two groups, the MMSE and Modified UPSIT scores are plotted versus age in Figure 3b. It can be seen that the two groups are entirely separable when modified UPSIT scores are used.

**Figure3.**
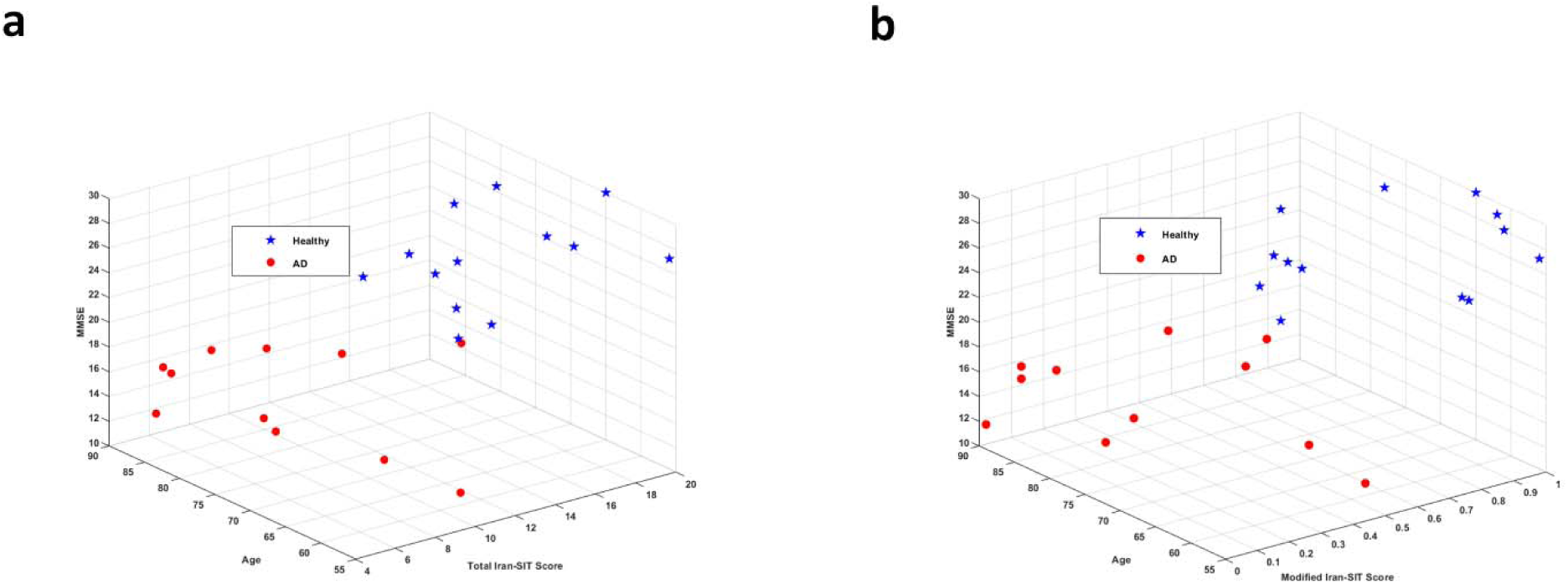
3D visualization of the MMSE score, total (a) and modified (b) UPSIT scores, and age.

To analyze which odors can discriminate between AD patients and healthy participants and which odors can still be perceived by AD patients, two visualizations of the UPSIT results are presented in Figure 4. The two significant odors (Grape and Chocolate) are indicated in both plots. In Figure 4a, the number of AD patients and healthy participants who answered each UPSIT odor identification question correctly is plotted. It can be seen that more than half of AD patients correctly identified the smells of Minty Toothpaste, Jasmine, Pineapple, and Strawberry. Figure 4b shows the UPSIT results when the participants in each group are divided into five-year age bins.

**Figure 4.**
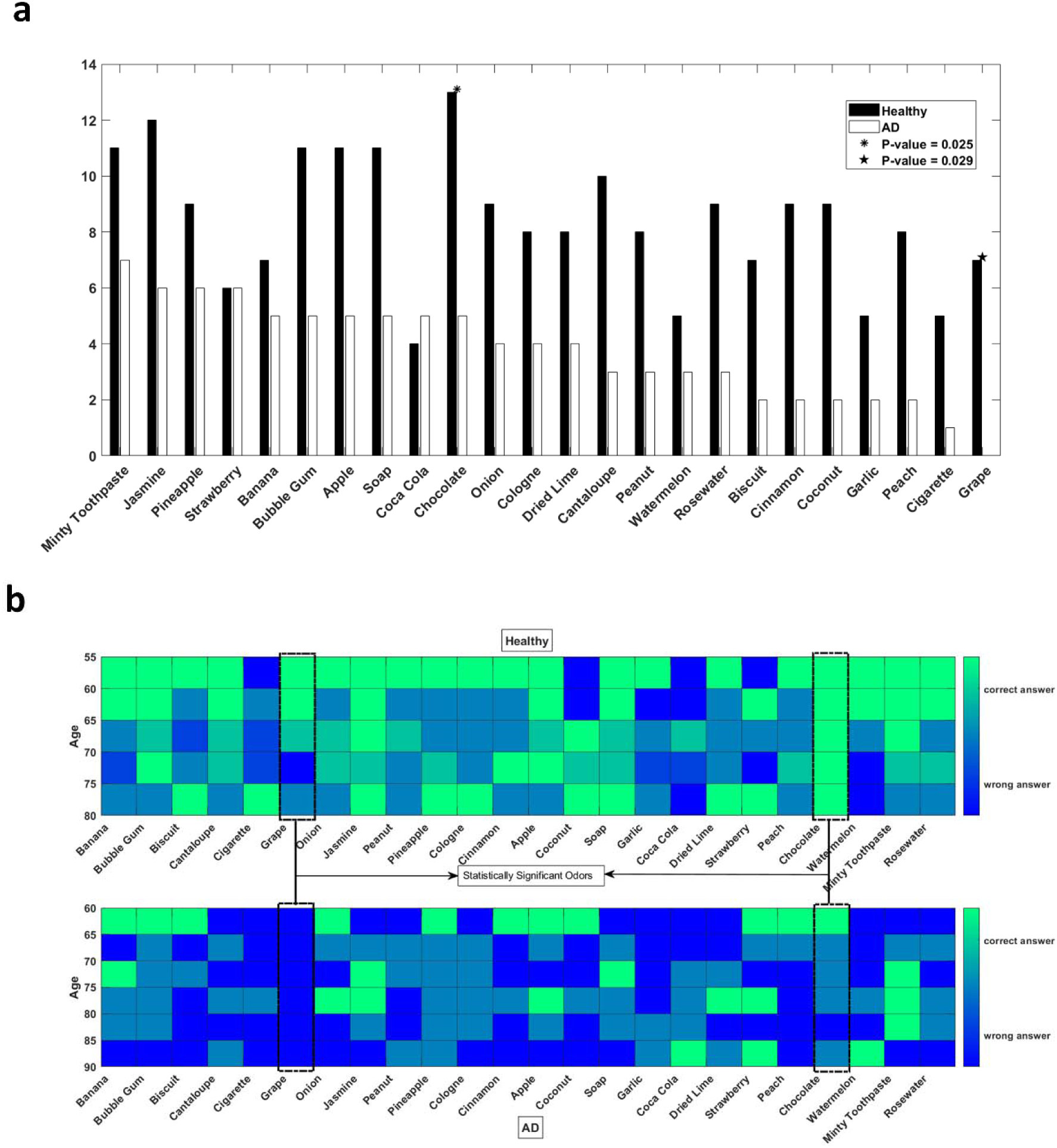
a) The number of correct answers for each UPSIT (Iran-SIT) odor identification question. The filled bars indicate the number of healthy participants, and the empty bars indicate the number of AD patients who answered each question correctly. The x-axis denotes the tested odors and is sorted from the left by the number of correct answers that the AD patients gave. Hence, the leftmost odor is the one that the AD patients identified most. The y-axis is the number of correct answers to each odor identification question. Two odors that are statistically significant in distinguishing between AD patients and healthy participants are denoted by an asterisk and a triangle. b) The number of correct answers for each UPSIT (Iran-SIT) odor identification question divided into five-year age bins. The shade of each bin denotes the number of correct answers. Green (light) pixels indicate that most participants in the corresponding age bin answered the question correctly, and the blue (dark) pixels suggest that most of the participants were unable to identify the questioned odor. The upper diagram is for the healthy participants, and the lower diagram is for the AD patients. The two statistically significant odors are denoted by dashed boxes.

### EEG Coherence

The imaginary part of coherence between each pair of EEG electrodes was calculated for each of the delta, theta, alpha, beta, and gamma frequency bands. To assess the significance of each of the six connections in the analysis, the mean value of the imaginary part of coherence across all five frequency bands was calculated for each connection. Statistical analysis indicated the imaginary part of coherence between the Fz and Cz channels to possess the highest significance in separating the two groups of AD patients and healthy participants. Figure 5a illustrates the relative significance of the six electrode-pair connections.

**Figure 5.**
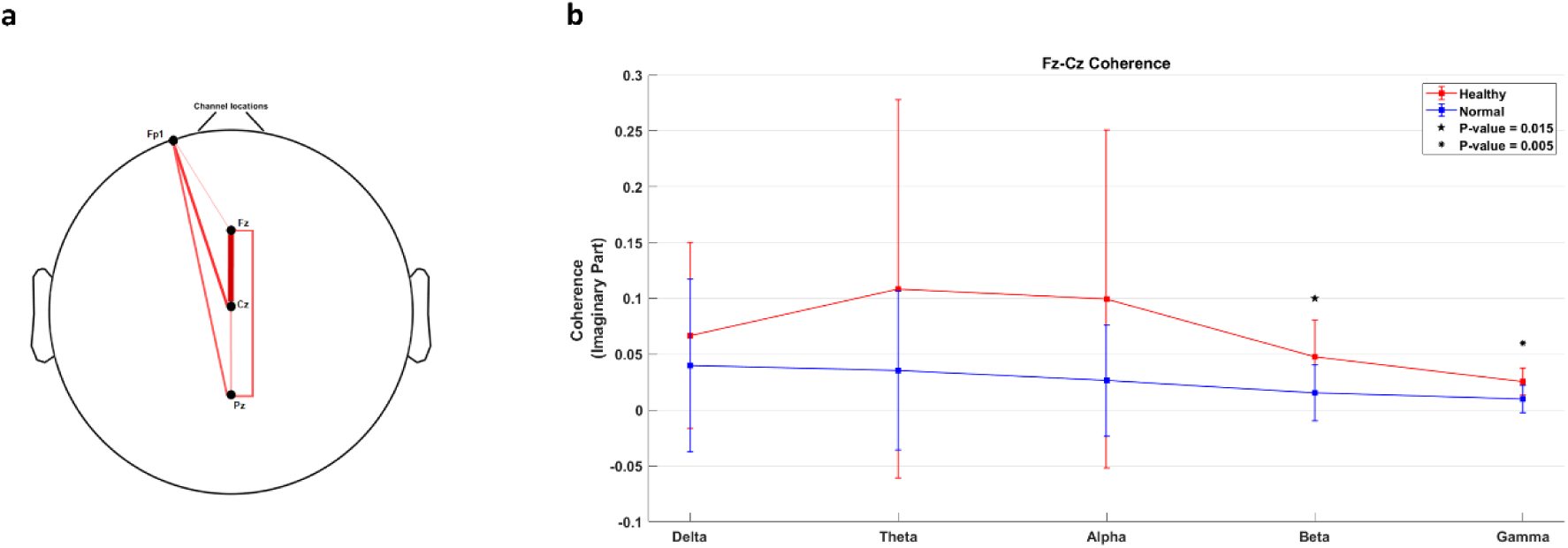
a) Difference in the imaginary part of coherence between the AD patients and healthy participants across the entire frequency range (0.5-40.5 Hz). Thicker lines indicate more statistically-significant connections. The Fz-Cz connection is the most significant connection with p-val<0.05. b) The imaginary part of coherence in the Fz-Cz connection measured in each frequency band. Statistically significant frequency bands are denoted by a star and a triangle. The blue line corresponds to healthy participants, and the red line corresponds to AD patients.

Then, p-values were calculated for the value of coherence in each frequency band for each connection, resulting in a total of 30 p-values (5 frequency bands times six connections). The Benjamini and Hochberg correction was applied using the effective sample size (38) calculated based on the correlation between the connections. This analysis resulted in the identification of the gamma (p-value = 0.005) and beta (p-value = 0.015) bands of the Fz-Cz connection to be significant. These two coherence values were selected as features for an SVM classifier of the two groups of participants.

We classified the EEG coherence data of the AD patients and healthy participants using a regressor combining the beta coherence, the gamma coherence and the age of participants. The results are shown in Table 4.

**Table 4.** Classification accuracy for different modalities and the multi-modal analysis based on significant components of each modality.

| Modality | Accuracy (%) | AUC |
| --- | --- | --- |
| Total UPSIT – Age | 83.3 % | 0.87 |
| Beta and Gamma Coherence in Fz-Cz - Age | 79.2 % | 0.85 |
| Beta and Gamma Coherence in Fz-Cz - Total UPSIT - Age | 83.3 % | 0.87 |
| **Beta and Gamma Coherence in Fz-Cz - Modified UPSIT - Age** | **91.7%** | **0.94** |

### Multimodal Analysis

The significant components identified in the behavioral olfactory test (UPSIT) and the EEG coherence analysis along with age were used as features to build an SVM classifier of AD patients and healthy participants. The performance of this multi-modal classifier, as well as its comparison with single-modality classifier results, are shown in Table 4.

The multi-modal classifier outperforms the single-modality classifiers in terms of accuracy. Besides, the resulting data collection protocol only requires the participants to answer two questions in the UPSIT test and the EEG data to be recorded from 3 electrodes (Fz, Cz, Fp1) and a reference (A1), offering a convenient procedure for examining elderly participants. The accuracy of the proposed multi-modal classifier is comparable to the MMSE-based classifier.

## Discussion

Standard diagnosis methods for AD based on CSF analysis involve invasive sampling and are hence not suitable for longitudinal studies (48). The proposed olfactory-based methodology in this paper is convenient to conduct and has considerably lower costs. It is also more accommodating for elderly patients as both the behavioral and EEG tests only require a small number of measurements. It hence provides a viable solution for monitoring the progress of the disease in a patient over time, and offers opportunities for longitudinal research studies.

In this study, we showed that the grape and chocolate odors could discriminate between AD patients and healthy participants with fair accuracy. Earlier studies have assessed the ability of different scents in similar tasks. For example, the study by Kjelvik et al. (2) identified some significant odors, which interestingly, chocolate was among them. A proposition for conducting olfactory-based studies would hence be to perform experiments in culturally-diverse groups of populations and identify the marker odors which are significant universally and those best suited for each culture or geography.

We also identified odors in our study which more than half of our AD patients could still perceive and successfully recognize. Understandably, a survey of a larger population is needed to identify such odors with more confidence. An interesting application for this set of scents would be to include them in olfactory assessment tests to discriminate between AD patients and individuals with anosmia or non-AD neurodegenerative diseases which can lead to the loss of olfactory functionality.

It should be noted that some odors could not be correctly identified by either of the healthy and AD groups. A reason for this lack of performance by the healthy participants might be the unfamiliarity of the study group with certain scents. For instance, both the AD and healthy groups were unable to identify the Coca-Cola odor because most Iranian elderly may have not experienced this smell in their daily life. These non-discriminating odors can be replaced in future sniffing kits by other scents to improve the performance of the test.

The result of EEG-based olfactory assessment suggested that the coherence in the gamma and beta bands significantly differ between AD patients and healthy participants. This result is in agreement with the evidence about the roles that the gamma and beta bands play in cognitive functions of the brain, and their deficit resulting from the neurodegenerative effects of AD. The high-frequency gamma oscillations (30-100 Hz) and the beta oscillations (13-30 Hz) appear to be particularly well suited for the maintenance of functions in the brain that involve *binding* the processed data from different sensory modules or elements stored in the memory (49). Multi-sensory data integration (50) (51), attentional sensory selection (52) (53) (54), working memory association (55), and generation of long-term memory through associations embedded as synaptic weight adaptation (56) (57), are all performed under the gamma and beta oscillatory regimes in the neuronal populations involved (49).

Employing high-frequency oscillations in functions that involve intricate recruitment of content from multiple sites in the brain is not a coincidental phenomenon; the resolution that high-frequency oscillations offer in their phase allows for fine-tuned coding of relative arrival times and latencies involved in accessing information and hence provides great input selectivity through high-precision control of spike timing (49) (58) (59).

Desynchronized neural activity and disruption of gamma oscillations have been observed in both human AD patients (60) (61) (62) (63) and the AD mouse models (64) (65) (66). As neuronal connectivity is affected by the accumulation of amyloid-β in the extracellular space (67), larger inhibitory circuits operating under high-frequency regimes turn into subpopulations that may produce these oscillations without synchrony with each other. In (68) the deficit in coherence between oscillations measured by EEG electrodes across the scalp in different frequency bands has been proposed as a diagnostic marker of dementia caused by Alzheimer’s disease.

While it is still a matter of debate whether these deficits in high-frequency oscillations are a consequence of the underlying disease progression or that they indeed play a causal role in inducing more biological changes that promote the disease (67), the deficit in the high-frequency oscillations can be associated on the functional level with the lowered *binding* activity in the cortex, causing the known symptoms of AD such as cognitive decline and dementia. Our study revealed the significance of the gamma and beta band coherences in separating AD patients and healthy participants and showed the difference to be more significant across the spatial range measured by the Fz and Cz electrodes. This is the scalp region close to the cortical areas known to be involved in many cognitive functions.

Our study showed that the olfactory deficit could be a fairly accurate marker for AD when behavioral assessment results are combined with the coherency results of the EEG recording. The accuracy of the proposed multi-modal classifier is significantly above the chance level (91.7%). Even if we develop a classifier based on the MMSE scores – which is based on tests that directly evaluate the participant’s memory and cognition – we may not always reach 100% accuracy. A common issue in performing the MMSE test is that some of its questions require reading and writing skills and therefore, illiterate subjects cannot get any scores from those parts. Also, running the test requires interaction between the participant and the memory specialist, increasing the probability of introducing bias during the test. Unlike MMSE, the proposed method in this study requires much less interaction and the behavioral olfactory assessment (UPSIT) test can be carried out even by the participants themselves if they have the ability to read the questions.

### Limitations

We have demonstrated the suitability of our methodology in identifying AD patients in a limited study on the Iranian population. The proposed method has to be applied to a bigger population in order to validate the usefulness of olfactory-based biomarkers in the diagnosis of AD. In addition, large-cohort research is needed to confirm the results, both in terms of validating the significance of the identified odors and coherence features, as well as examining the outcome when patients suffering from non-AD dementia or moderate and severe AD patients are also present in the study.

As odor perception is a culture-dependent phenomenon, the exact results derived in our study may not be directly applicable to different populations. A more comprehensive study comprising of populations of participants from different cultural backgrounds is needed to verify the validity of the proposed approach in general, and the significance of the individual scents or other features used in our regressors and classifiers in particular.

A related notable remark is that while the results of the smell identification test (UPSIT) may have strong cultural dependencies, the EEG-based coherence analysis may prove to be a relatively more robust procedure. This is due to the fact that higher-level functions of the brain are represented in the coherence values measured in the EEG analysis, and some level of abstraction from the particular smells that are perceived may be represented in these measurements.

Another limitation of the smell identification test is that successfully answering questions in it involves both the perception of the presented odor as well as its recognition through a memory recall process. This indicates an inherent ambiguity in this test between a lack of perception of the presented smell and failure to identify the name of the odor which may have indeed been perceived. The EEG-based olfactory assessment is advantageous in this aspect to the UPSIT test as it mainly focuses on the perception ability and not the identification or naming functions.

### Extensions

One interesting extension of this work is to repeat the EEG-based experiments using the two odors (chocolate and grape) which were identified in the olfactory recognition task as significant and compare the results of the coherence analysis with the current results. As these two odors best separate the two groups of participants, it is interesting to see any gains their usage may provide to the single-modal EEG-based coherence results.

An essential extension to the EEG analysis is to examine other approaches, such as the difference between the responses to the two odorants within each group of AD patients and healthy participants. There are two possible ways to derive these differences. One is to repeat the current coherency-based approach separately for each of the odorants. The other is to make the comparison directly in the temporal domain of the recorded EEG data after artifact and noise removal. Possible advantages of these odorant-differential analyses may include the additional dimension that they provide as the difference in responses to the two presented odors. To perform this extension, it is better to conduct the experiments with a sequence of odors in which both the odorants are presented randomly with equal probability so the number of epochs related to each odorant would be comparable.

Another possible domain for extending the EEG-based olfactory test is to repeat the experiment for each participant in more than one session and use different pairs of odorants in each experiment. This allows for studying the sensitivity map of each participant relative to different odors. Running an experiment with more than two odorants presented by the same olfactometer is a possibility in this domain, but requires modifications in the design of the olfactometer.

On the cohort design side, a necessary extension is to recall the participants for another round of experiments after a period of six to twelve months and perform longitudinal analyses to study the correlation between the olfactory decline and the progress of the AD in each patient. Another critical extension that can further evaluate the specificity power of the proposed approach is to include a third participant group consisting of non-AD MCI patients and examine the single- and multi-modal analyses of the current study in discriminating the three groups from one another. These two latter extensions are within the scope of our on-going data collection campaign. Any additional results achieved in each of these extended studies will be provided in future reports.

## Conclusions

In this paper, we have demonstrated the efficacy of an inexpensive methodology for evaluating the olfactory deficit in the elderly population for being utilized as a marker of AD with fair accuracy. Our proposed approach combines behavioral olfactory data with EEG measurements to yield an accurate assessment of the participant’s state.

Statistical analysis of the results of the smell identification test yields two odors (Chocolate and Grape) as significant (p-values < 0.05) from a set of 24 odorants. The EEG coherence analysis indicates the gamma and beta bands to be significant (p-values < 0.05) in the link between the Cz and Fz channels, with the gamma band possessing a much higher significance. The proposed multi-modal classifier yields an accuracy of 91.7% in separating AD patients from healthy participants.

The accessibility and low cost of the proposed procedure allow for large-scale screening of AD in different geographical regions, a looming necessity across the world as the aging population is rapidly expanding. Furthermore, the results of this work can provide researchers with new insights about the relationship between AD progression and olfactory deficit and can lead to new treatment methods based on olfactory stimulation.

## Acknowledgments

The authors would like to thank Ziaeian Hospital in Tehran for providing staff time and equipment for data collection in this study. We are grateful to the patients and their families who participated in this study.

