## supplementary materials for "Olfactory Response as a Marker for Alzheimer’s Disease: Evidence from Perception and Frontal Oscillation Coherence Deficit"

**S1. Iran-SIT Test Questions**

A list of Iran-SIT odors is shown in the table below (the correct answers, i.e. the presented odors are denoted in boldface).

|  | A | B | C | D |
| --- | --- | --- | --- | --- |
| 1 | Gasoline | Cherry | **Banana** | Garlic |
| 2 | Pizza | Cucumber | Alcohol | **Bubble Gum** |
| 3 | Cologne | **Biscuit** | Tuberose | Orange |
| 4 | Cinnamon | Vinegar | **Cantaloupe** | Kebab |
| 5 | Peach | Saffron | **Cigarette** | Tangerine |
| 6 | **Grape** | Roasted Seed | Parsley | Fried Chicken |
| 7 | Melon | Rose | Coffee | **Onion** |
| 8 | Lemon | **Jasmine** | Smoke | Fish |
| 9 | **Peanut** | Celery | Apple | Cardamom |
| 10 | Tea | Tobacco | **Pineapple** | Gas |
| 11 | Cake | Sour Orange | Black Pepper | **Cologne** |
| 12 | **Cinnamon** | Pomegranate | Egg | Washing Machine Powder |
| 13 | Olive | Sewage | **Apple** | Butter |
| 14 | **Coconut** | Rosewater | Sausage | Smoke |
| 15 | Gourmeh Sabzi (Persian Herb Stew) | **Soap** | Lemon | Watermelon |
| 16 | Chocolate | Saffron | Cucumber | **Garlic** |
| 17 | Basil | Pear | Fish | **Coca Cola** |
| 18 | Hami Melon | **Dried Lime** | Honey | Grapes |
| 19 | Olive | **Strawberry** | Leather | Thyme |
| 20 | Peanut | **Peach** | Coffee | Pizza |
| 21 | **Chocolate** | Sour Orange | Vinegar | Sewage |
| 22 | Garlic | Cigarette | **Watermelon** | Bread |
| 23 | Rice | Pistachio | Pepper | **Minty Toothpaste** |
| 24 | **Rosewater** | Orange | Gasoline | Cheese |

**S2. Plot of MMSE Scores versus Age**

Healthy participants are denoted by blue asterisks and AD patients are indicated by red circles. A regression line is fitted to the data of each group, and the confidence bounds are also plotted.

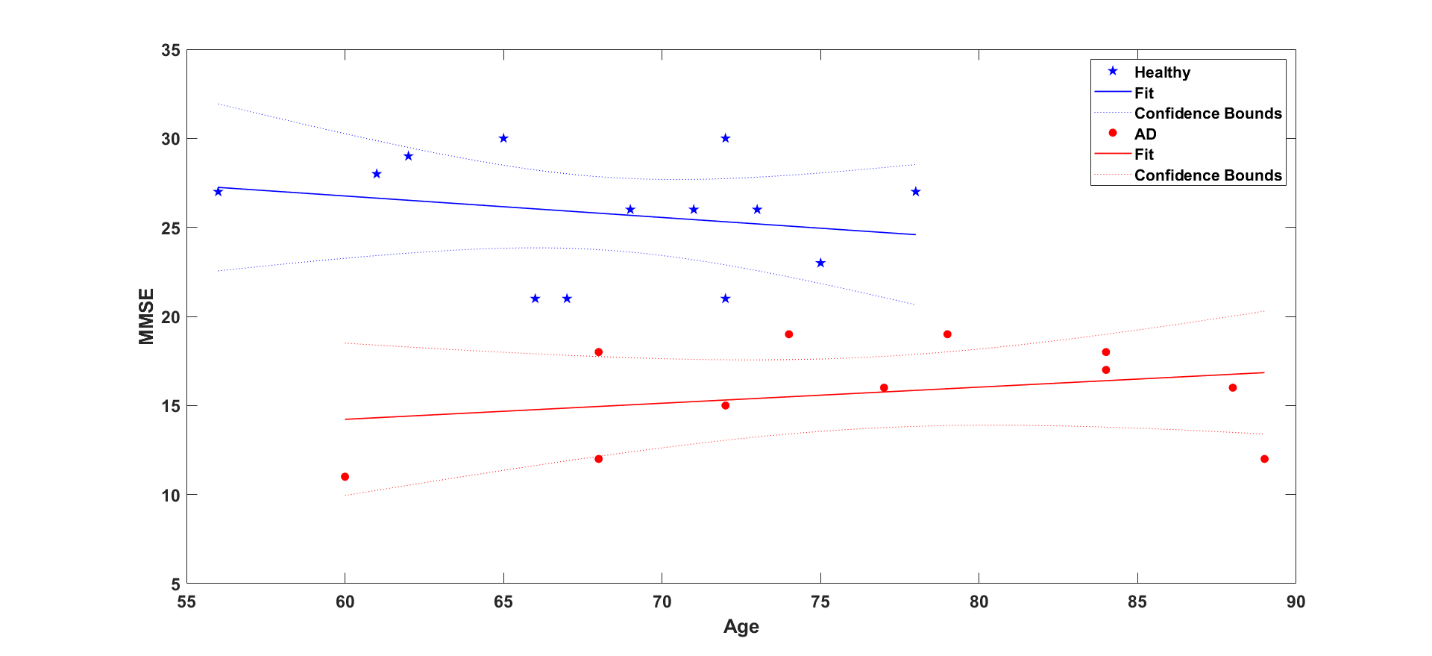

**S3. EEG Channel Locations**

EEG data were recorded from the Fp1, Fz, Cz, and Pz electrodes referenced to earlobe. Channel locations are shown in the figure below.

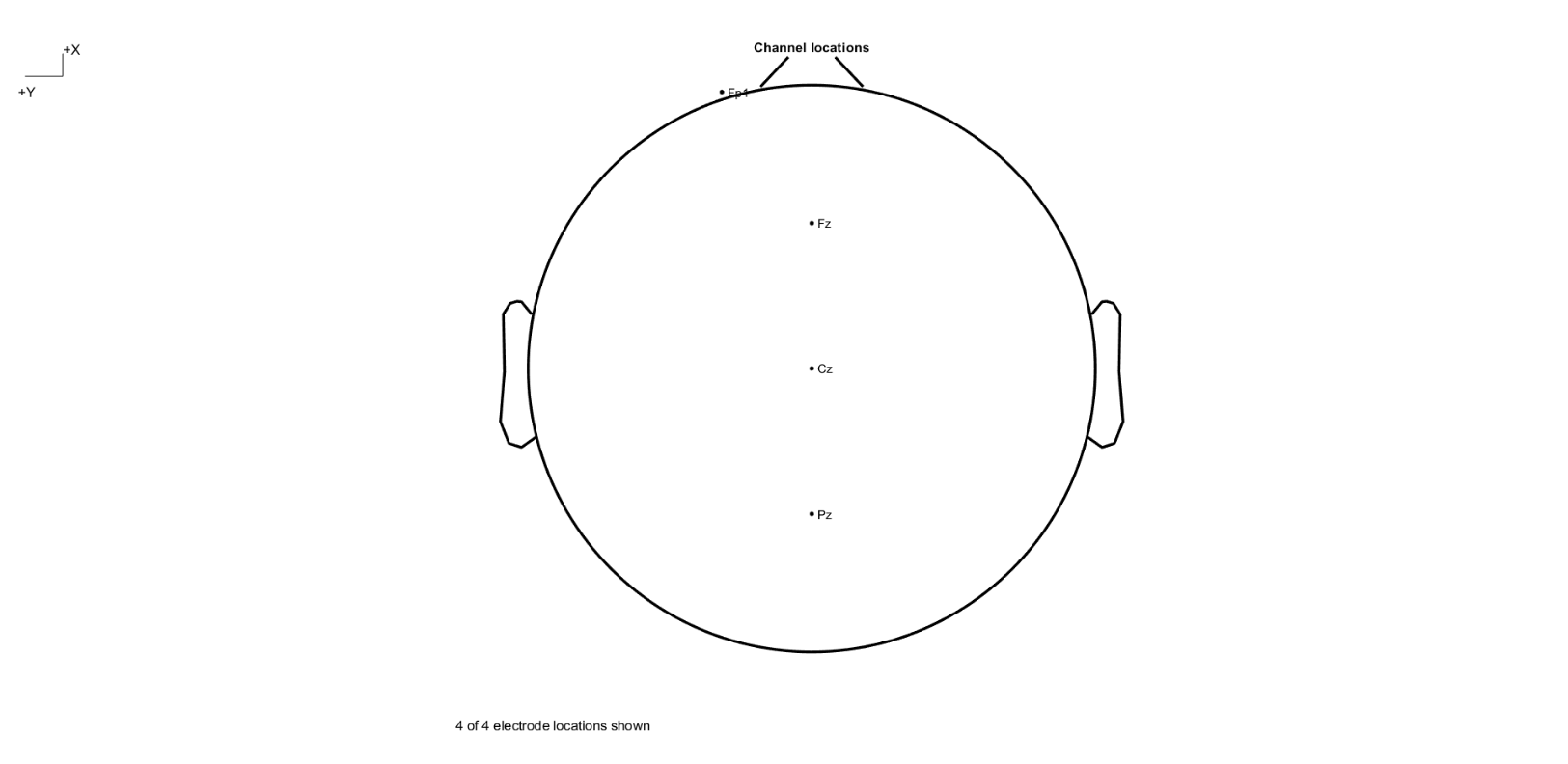

**S4. Plot of the imaginary part of coherence versus age in the Fz-Cz connection in a) beta band, b) gamma band**

Data of AD patients are denoted by red circles and of healthy participants by blue asterisks. A line is fitted to the data of each group. A higher value for coherence implies weaker connectivity. Except for the gamma band in the AD data, the increase in coherence (decline in connectivity) has a small correlation with age.

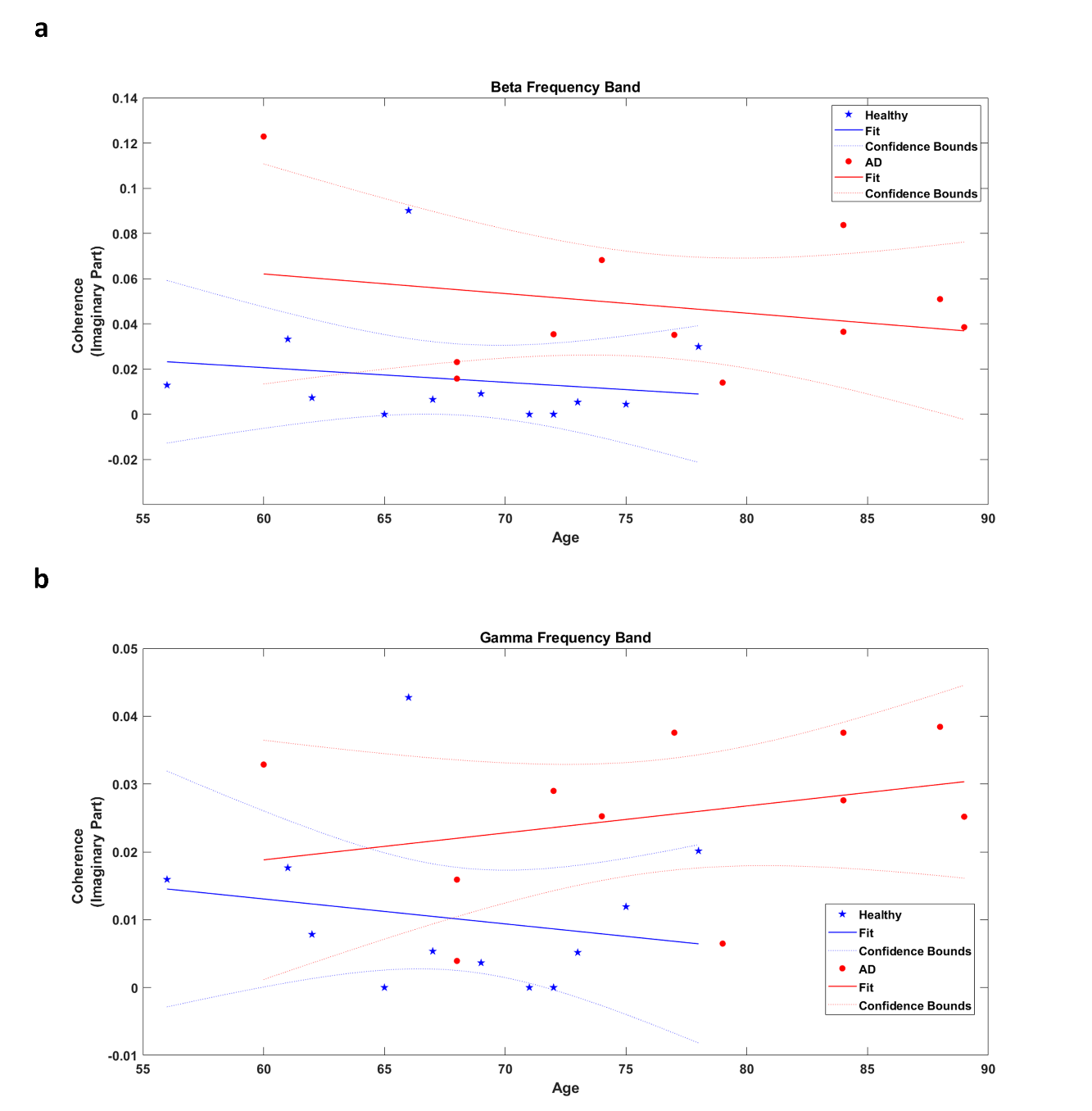

**S5. Plot of Total UPSIT Score versus Total MMSE Score**

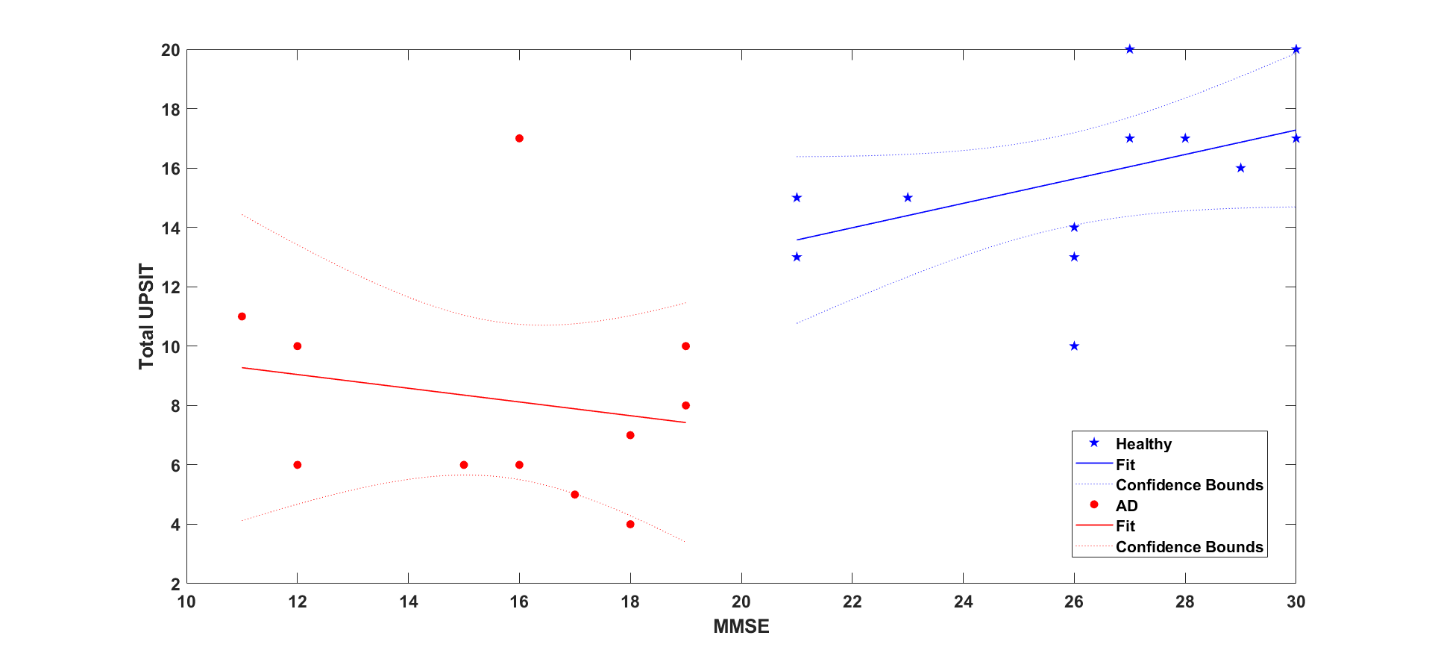
